# Bibliometric Analysis of Circular RNA Cancer Vaccines and Their Emerging Impact

**DOI:** 10.1101/2024.08.31.610618

**Authors:** Uddalak Das, Soupayan Banerjee, Meghna Sarkar

## Abstract

Circular RNA (circRNA) vaccines are emerging as a revolutionary strategy in cancer immunotherapy, offering novel mechanisms for inducing robust and durable immune responses. Unlike traditional linear mRNA vaccines, circRNAs exhibit exceptional stability, enhanced translational efficiency, and resistance to exonuclease degradation, making them ideal candidates for vaccine development. Recent studies have shown that circRNA vaccines play an important and specialized role in cancer treatment. However, there are currently no relevant bibliometric studies. This study aimed to apply bibliometrics and scientometrics to summarize the knowledge structure and research hotspots regarding the role of circRNA vaccines in cancer. Publications related to circRNA vaccines in cancer were searched on the Scopus database. VOSviewer and Scopus were used to conduct the analyses. This study summarizes the research trends and development of circRNA vaccines for cancer and also a comparative analysis between mRNA cancer vaccine and circRNA cancer vaccine identifying potential areas of focus, innovation and growth. This information will provide a reference for researchers to studying circRNA vaccines against cancer due to its increasing trends over recent times.

## 1. INTRODUCTION

Cancer remains a global threat with high incidence and mortality rates. Recent advancements in cancer immunotherapy offer a promising alternative to conventional treatments like chemotherapy, surgery, and radiation therapy Siegel (Siegel *et al*., 2023). Unlike these traditional methods, which solely target cancer cells, immunotherapy harnesses the body’s immune system to combat cancer by activating anti-tumor responses and modifying the tumor microenvironment, leading to tumor shrinkage and improved patient survival.

Despite progress in cancer treatment, conventional therapies like chemotherapy, radiation, and immune checkpoint inhibitors (ICIs) face challenges, including limited effectiveness, adverse effects, and treatment resistance. These limitations underscore the need for innovative strategies that stimulate durable antitumor immune responses. While traditional therapies effectively target rapidly dividing cancer cells, they often cause significant side effects and may not provide lasting results, leading to tumor recurrence and resistance (Labani-Motlagh *et al*., 2020). Moreover, while checkpoint inhibitors targeting PD-1 or CTLA-4 have shown success in some cancers, their efficacy is limited to certain patients, with resistance arising due to changes in the tumor microenvironment and immune escape mechanisms (Shiravand *et al*., 2022).

Addressing these challenges, personalized medicine has emerged, focusing on patient-specific neoantigens. Technological advances like whole-genome mapping and next-generation sequencing (NGS) have facilitated the development of neoantigen vaccines, offering highly immunogenic epitopes for enhanced immune priming. An ideal immunotherapeutic cancer vaccine should trigger specific immune responses against tumor-specific antigens (TSAs), maintain long-term tumor memory, and minimize adverse effects. However, translating this ideal vaccine into practical therapies has been challenging. To overcome these hurdles, researchers are exploring ways to increase tumor antigen immunogenicity and counteract tumor immunosuppression.

CircRNA-based cancer vaccines offer a promising solution, leveraging the stability and extended antigen expression of their covalently closed-loop structures. Unlike linear mRNA vaccines, circRNAs resist RNase degradation, enabling sustained immune activation and improved therapeutic efficacy (Zlotorynski, 2019). Their ability to encode TSAs and adjuvants allows for customizable vaccine designs tailored to individual tumor profiles, addressing tumor heterogeneity (Buonaguro & Tagliamonte, 2020).

CircRNA-based vaccines also offer additional benefits in cancer treatment. Their stability and prolonged antigen expression enhance immune responses, potentially overcoming immune evasion mechanisms (Niu *et al*., 2023). Moreover, circRNAs can function as both immunogens and adjuvants, simplifying vaccine formulation and boosting immunogenicity. This flexibility allows for the incorporation of multiple TSAs and adjuvants in a single structure, maximizing the immune response against cancer cells (Amaya *et al*., 2023). CircRNA vaccines also demonstrate scalability and cost-efficiency in production, with their simple composition and durability facilitating large-scale manufacturing and storage, making them more accessible and affordable compared to complex biological products. Furthermore, circRNA vaccines promise precise administration and improved tissue penetration, reducing side effects and optimizing therapeutic outcomes (Xie *et al*., 2023).

In an era where the rapid expansion of information and big data technologies profoundly influences all fields, including healthcare and medical research, the relevance of systematic bibliometric studies has never been greater (Schlick *et al*., 2018; Trifirò *et al*., 2018). Bibliometrics uses mathematical and statistical methods to analyze research trends and achievements in specific fields, helping scholars quickly identify current hotspots and guide future research directions. Tools like CiteSpace (Chen, 2006), VOSviewer (Van Eck & Waltman, 2010), and the R package ‘bibliometric’ are commonly used to visualize scientrometric data across various fields, including cancer, surgery, and genetics. However, there is a notable lack of bibliometric studies specifically focused on circular RNA based vaccines for cancer. While Luofei Zhang conducted a bibliometric analysis on vaccines and cancer prevention (Zhang *et al*., 2023), it did not cover RNA vaccines. Yang *et al*. (2023) conducted a bibliometric analysis of RNA vaccines for cancer but did not focus on the emergence of circRNA based cancer vaccines. This highlights the pressing need for a comprehensive bibliometric analysis of recent circRNA-based cancer vaccines to reshape the landscape of cancer immunotherapy, guiding future research and innovation in this critical area. Given the potential of circRNA vaccines in cancer treatment, we conducted a bibliometric analysis of literature on circular RNA vaccines in cancer immunotherapy. Additionally, we performed a comparative analysis to identify key research trends, gaps, and future directions in this emerging field. Our study aims to fill the existing void in bibliometric research on circRNA-based cancer vaccines, providing a foundational framework to guide future investigations and accelerate advancements in cancer immunotherapy.

## 2. METHODOLOGY

### 2.1. Search strategy

The literature search was conducted to generate inclusion and search terms based on the population, idea, and content requirements as well as the review objectives. The search phrases and words from the free text were joined together using the Boolean operators “AND” and “OR.” The search keywords were “RNA AND cancer AND vaccine OR mRNA AND cancer AND vaccine” for RNA cancer vaccines and “circRNA AND cancer AND vaccine OR circular AND RNA AND cancer AND vaccine” for circRNA based cancer vaccines, limited within title/abstract/keywords. Electronic databases (Scopus and PubMed) and manual methods (searching reference lists of included publications) were used for the literature searches. Publications written in English that were released between 2013 and 2024 are acceptable for consideration.

The initial search identified 5,270 articles for RNA vaccine and 106 articles for circRNA based cancer vaccines. To refine the selection, only English-language papers were considered, reflecting the predominant use of English in academic discourse. Subsequently, two categories of literature -research articles, and review papers were selected. Consequently, a total of 4,550 publications were retained for RNA cancer vaccines and 96 publications were retained for circ-RNA cancer vaccines for further analysis.

### 2.2. Data Analysis

Bibliometric data were sourced from the Scopus database, a widely recognized platform for bibliometric analysis that provides extensive information on scholarly publications. A search was performed in Scopus on August 13, 2024, covering the period from 2013 to 2023. The bibliometric analysis was partially conducted using VOSviewer (version 1.6.2), a specialized software for analyzing key information from a large volume of publications, which was utilized to construct collaboration, co-citation, and co-occurrence networks. Full counting method, giving same weightage to each co-authorship, co-occurrence, bibliographic coupling, or co-citation link. Additionally, Microsoft Office Excel 2019 was employed for various analyses of the publication data.

## 3. RESULTS AND DISCUSSION

### 3.1. Comparative Analysis of RNA and circRNA-based Vaccines for Cancer

#### 3.1.1. Quantitative analysis of publication

The comparative analysis of research trends in “circRNA-based vaccines on cancer” and “RNA-based vaccines on cancer” from 2013 to 2024 shows distinct patterns (Figure 1). For circRNA-based vaccines (Panel A), there was a slow growth from 2013 to 2020 with a noticeable surge peaking in 2023, followed by ongoing works in 2024. In contrast, RNA-based vaccines (Panel B) exhibited stable publication numbers from 2013 to 2019, a significant rise starting in 2020, peaking in 2022, and a decrease from 2023.

**Figure 1.**
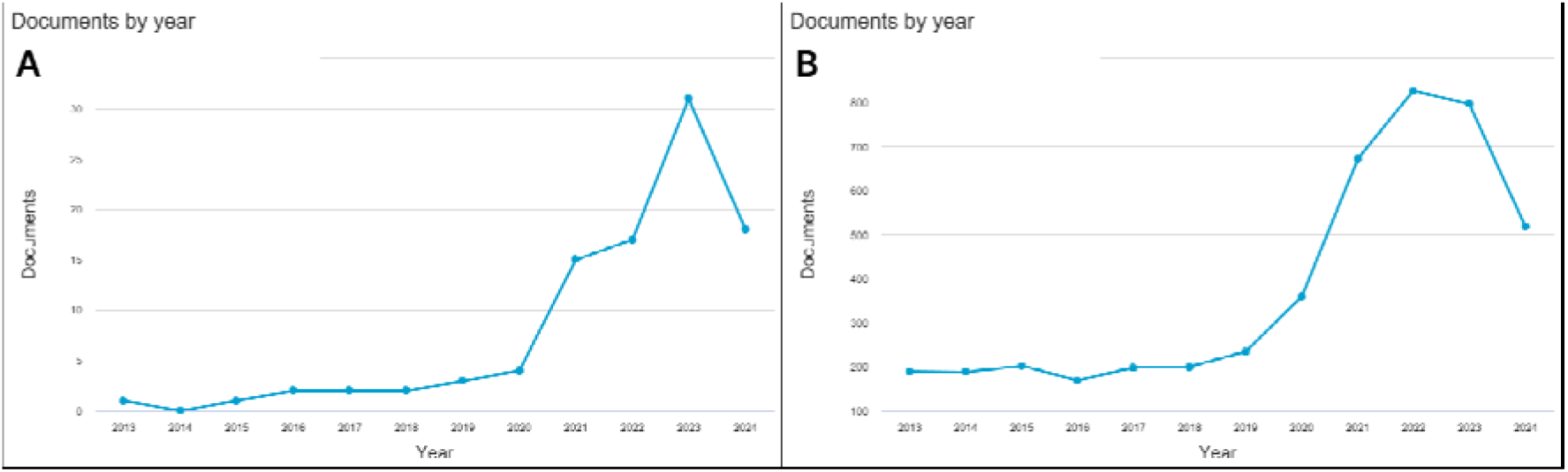
Annual output of articles A: Circular RNA cancer vaccines B: RNA Cancer vaccines

The data reveal that RNA-based vaccines have a longer research history and saw a boost due to advancements in RNA technology, particularly with COVID-19 vaccines. Conversely, circRNA-based vaccines are a newer focus with rapid growth post-2020, reflecting increased interest in their unique advantages such as stability and potential for innovative cancer treatments. The decline in both fields in 2024 may be due to publication lag or shifting research focuses. The emerging prominence of circRNA-based vaccines suggests they are becoming a critical area of research with potential to significantly impact cancer immunotherapy.

#### 3.1.2. Authors

The analysis of contributions and collaborations in circRNA-based vaccines on cancer (Figure 2: Panels A and B) and RNA-based vaccines on cancer (Figure 2: Panels C and D) reveals distinct patterns. Panel A shows that the top 15 authors in circRNA-based vaccine research have a uniform distribution of publications, indicating a nascent field with no dominant leaders. Panel B’s co-authorship network for circRNA-based vaccines is sparse, with only seven out of 25 authors showing interconnected collaborations. In contrast, Panel C reveals that the top 15 authors in RNA-based vaccine research have a more diverse and concentrated publication output, reflecting a mature field with prominent researchers. Panel D shows a well-established co-authorship network in RNA-based vaccine research, with 827 out of 1262 authors exhibiting extensive interconnected collaborations.

**Figure 2.**
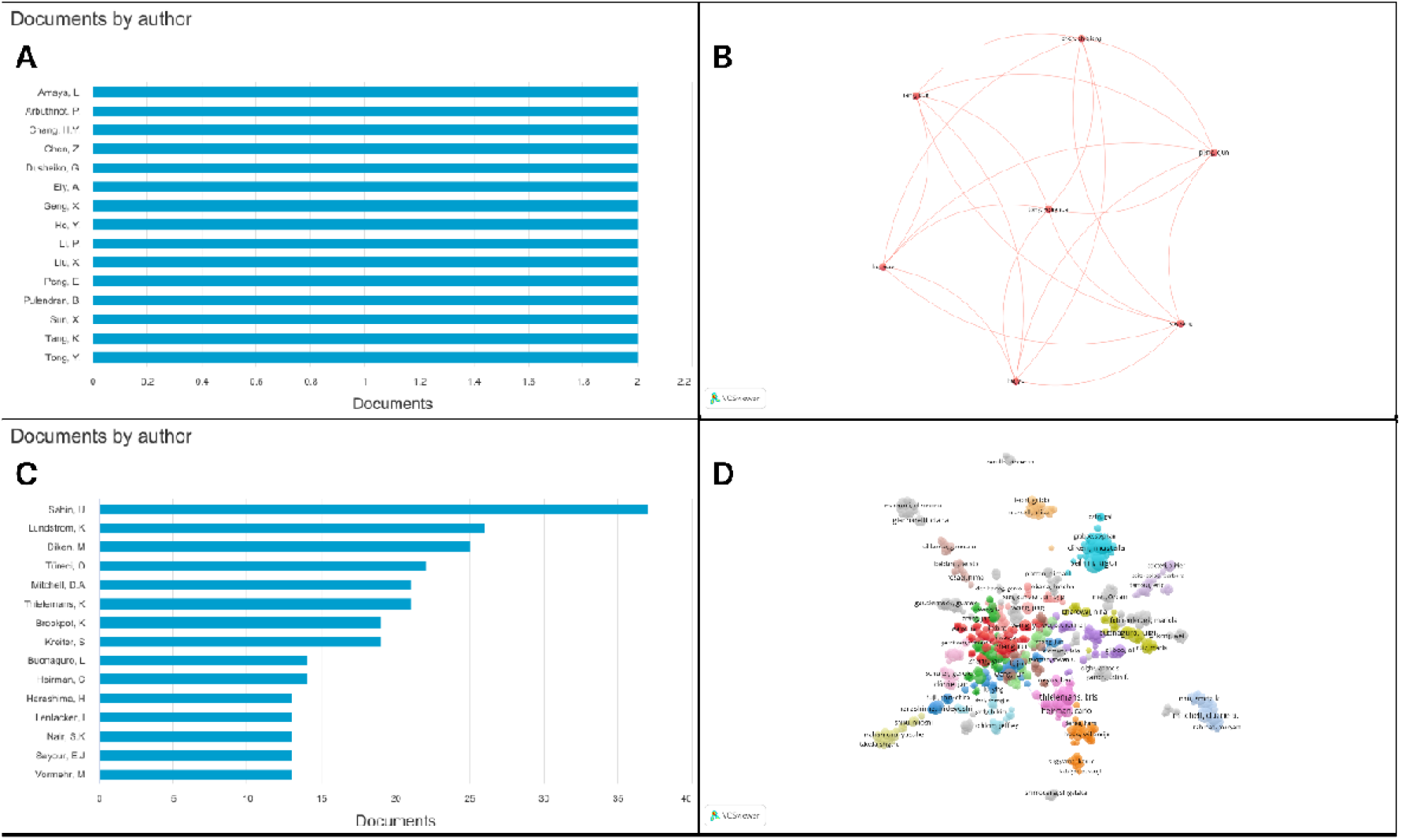
Author contributions A: Top 15 authors In circular RNA cancer vaccines B: Co-authorship network of circular RNA cancer vaccines C: Top 15 authors in RNA Cancer vaccines D: Co-authorship network of RNA cancer vaccines

The data indicate that circRNA-based vaccines are still in the early stages of development, as evidenced by the uniform author contributions and sparse co-authorship networks in Panels A and B. This suggests that foundational research is ongoing with emerging collaboration. Conversely, RNA-based vaccines are characterized by a mature and prolific research community, with well-established authors and dense collaborative networks, as shown in Panels C and D. The contrasting results highlight that while RNA-based vaccines benefit from extensive collaboration and experience, circRNA-based vaccines are poised for growth. The emerging nature of circRNA-based vaccines implies potential for increased collaboration and productivity, similar to the trajectory of RNA-based vaccines.

#### 3.1.3. Institutional Analysis

The analysis of institutional contributions and collaborations from 2013 to 2024 reveals distinct patterns in circRNA-based and RNA-based vaccines on cancer (Figure 3). Panel A shows that the top 15 institutions in circRNA-based vaccine research have a narrow range of document outputs, with none exceeding eight publications, indicating the field’s early stage. Panel B highlights a sparse network with only four out of 354 institutions meeting the minimum threshold of two documents, and no interconnected collaborations. In contrast, Panel C shows that the top 15 institutions in RNA-based vaccine research have significantly higher publication counts, with some exceeding 100 documents, reflecting a more mature field. Panel D demonstrates a robust network of 47 interconnected institutions out of 238 meeting the threshold of three documents, showcasing an advanced stage of collaboration and research activity.

**Figure 3.**
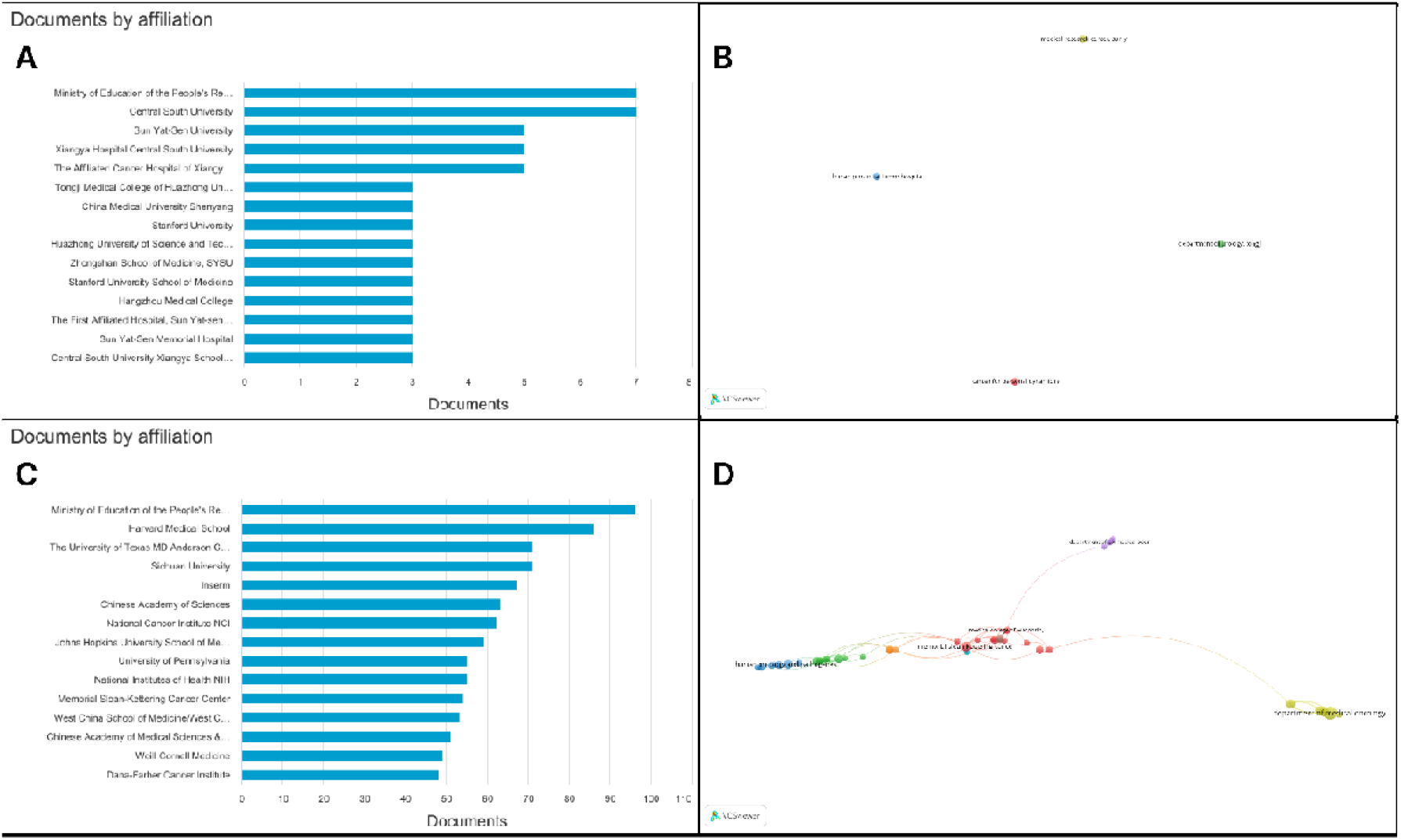
Affiliated universities/organization/institutions A: Top 15 organizations In circular RNA cancer vaccines B: Organizations network of circular RNA cancer vaccines C: Top 15 organizations in RNA Cancer vaccines D: Organizations network of RNA cancer vaccines

The results indicate that circRNA-based vaccines are still in their infancy, as evidenced by the limited publication output and sparse collaborative networks in Panels A and B. This suggests that the field is in the exploratory phase, with research efforts dispersed and foundational work still being established. On the other hand, RNA-based vaccines have a well-developed research environment with high publication output and extensive collaborative networks, as shown in Panels C and D. The advanced stage of RNA-based vaccine research highlights a mature, competitive field with established institutions driving innovation. The emerging nature of circRNA-based vaccines presents significant opportunities for growth, potentially following a trajectory similar to RNA-based vaccines as the field develops and institutional engagement increases.

#### 3.1.4. Funding Agencies

In figure 4, panel A reveals that the National Natural Science Foundation of China (NSFC) is the leading funder for circRNA-based cancer vaccines, supporting approximately 30-35 documents, with the Ministry of Science and Technology of China also contributing. This suggests that circRNA-based vaccine research is emerging but still relatively limited, with primary funding concentrated in China. In contrast, Panel B shows a more diverse funding landscape for RNA-based cancer vaccines. The National Institutes of Health (NIH) is the top funder, supporting over 500 documents, followed by the NSFC. The presence of European institutions such as the European Commission and Deutsche Forschungsgemeinschaft (DFG), as well as other major agencies like the National Cancer Institute and the Japan Society for the Promotion of Science, highlights the global and mature nature of RNA-based vaccine research.

**Figure 4.**
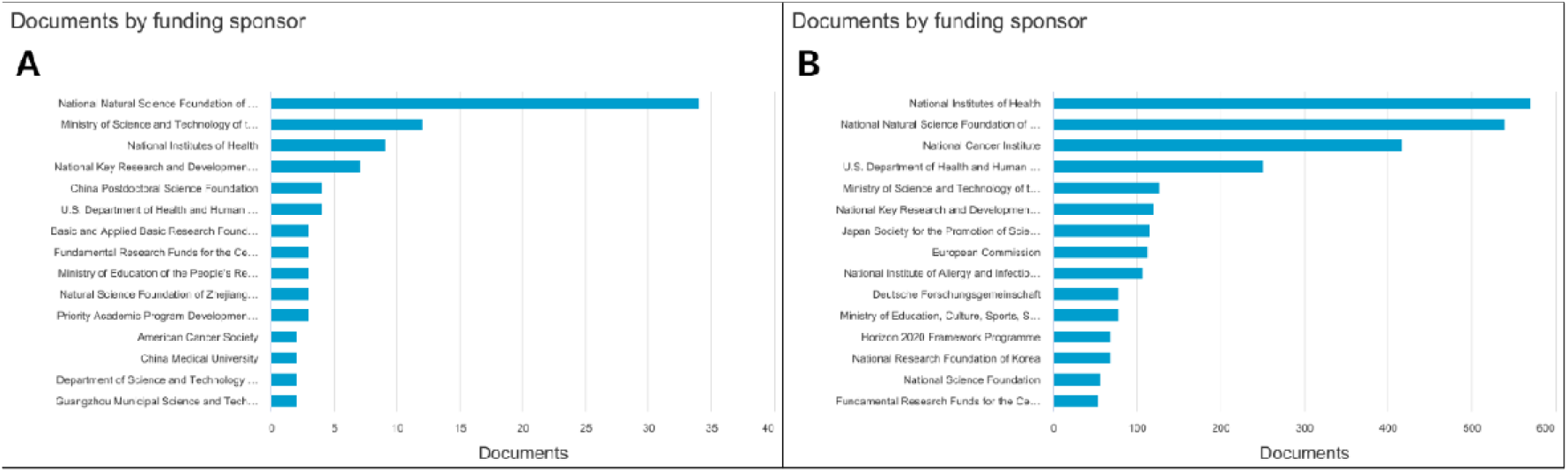
Top 15 Funding organizations A: Circular RNA based cancer vaccines B: RNA based cancer vaccines

The results clearly show that circRNA-based vaccines are in the early stages of research with limited funding and fewer documents per institution, primarily concentrated in China (Panel A). The NSFC’s leading role reflects a focused but developing interest in this area. On the other hand, RNA-based vaccines benefit from extensive global funding and a broad range of supporting institutions (Panel B), reflecting a mature and well-established field with high publication volumes and significant international collaboration. The disparity in funding scale highlights the nascent stage of circRNA-based vaccines compared to the well-developed RNA-based vaccine sector. However, increasing interest from major global funders, including the NIH and other U.S. and European institutions, suggests a promising future for circRNA-based vaccines, with potential for rapid growth as the field gains traction and research advances.

#### 3.1.5. Countries

In figure 5, panel A shows that China and the United States are the top contributors to circRNA-based cancer vaccine research, each with over 50 documents. Other countries like Germany, India, and Iran have significantly lower outputs. This indicates that circRNA-based vaccine research is still concentrated in a few leading countries and has not yet achieved global prominence. In Panel C, the United States leads RNA-based cancer vaccine research with over 1600 documents, followed by China, Germany, and Italy. This widespread international involvement reflects the mature and established nature of RNA-based vaccine research, with significant contributions from numerous countries, including Japan, France, and Canada. Panel B’s network analysis for circRNA-based vaccines reveals that out of 34 involved countries, only 13 meet the minimum threshold for interconnections, and only 9 show meaningful connections, with China central to the network. This sparse connectivity highlights the early-stage and localized nature of circRNA research. In contrast, Panel D shows a highly interconnected network for RNA-based vaccines, with 176 countries meeting the threshold and a dense web of interactions centered around the United States, China, Germany, and Japan. This indicates a well-established and globally integrated research community.

**Figure 5.**
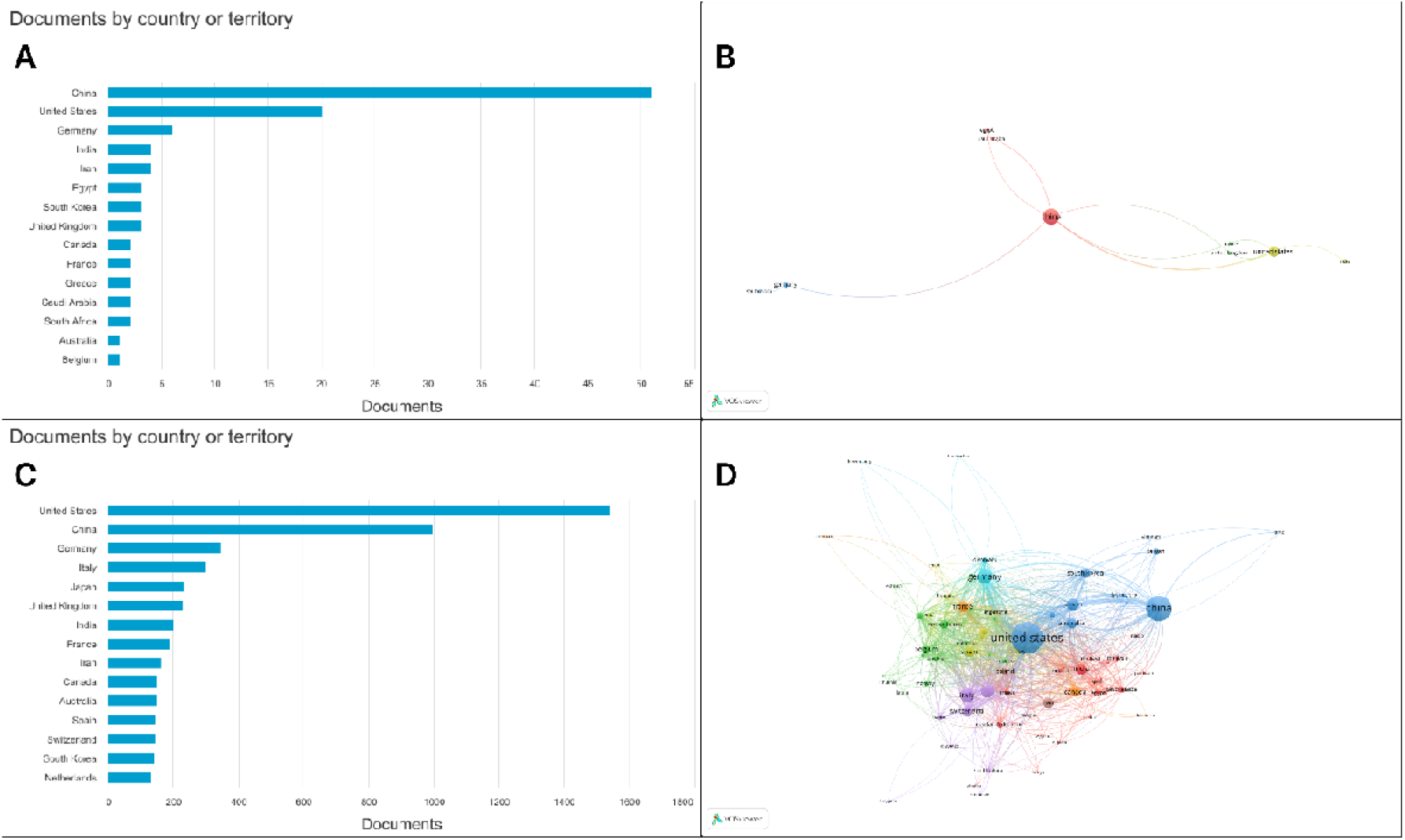
Figure 3 Countries A: Top 15 countries In circular RNA cancer vaccines B: Countries network of circular RNA cancer vaccines C: Top 15 countries in RNA Cancer vaccines D: Countries network of RNA cancer vaccines

The results highlight that circRNA-based vaccine research is currently limited to a few key countries (China and the US) with relatively few international collaborations (Panel A and B). This contrasts sharply with the broader, more mature global network of RNA-based vaccine research (Panel C and D). The significant disparity in document numbers and network complexity reflects the advanced state and extensive global collaboration in RNA-based vaccine research, driven by successful applications like mRNA vaccines for COVID-19. The circRNA field, while promising due to its potential advantages, such as stability and durable immune responses, is still developing. Increased international collaboration and funding are crucial for circRNA-based vaccines to achieve a similar level of global integration and research maturity as seen in RNA-based vaccines. The current concentration in China and the US, coupled with limited global connectivity, suggests an opportunity for growth and broader international engagement in circRNA vaccine research in the future.

#### 3.1.6. Major Subject Areas

As shown in figure 6, panel A reveals that circRNA-based vaccine research is predominantly focused on Medicine (30.7%), Biochemistry, Genetics, and Molecular Biology (28.1%), and Pharmacology, Toxicology, and Pharmaceutics (15.6%). This highlights a strong emphasis on clinical applications and the biological mechanisms of circRNA, with a notable focus on preclinical drug development. Panel B for RNA-based vaccines shows a broader distribution, with significant attention in Medicine (32.1%), Biochemistry, Genetics, and Molecular Biology (23.6%), and increased focus on Immunology and Microbiology (14.8%). It also includes contributions from Chemical Engineering, Chemistry, and Materials Science, reflecting a mature, interdisciplinary approach crucial for developing vaccine delivery systems and formulations.

**Figure 6.**
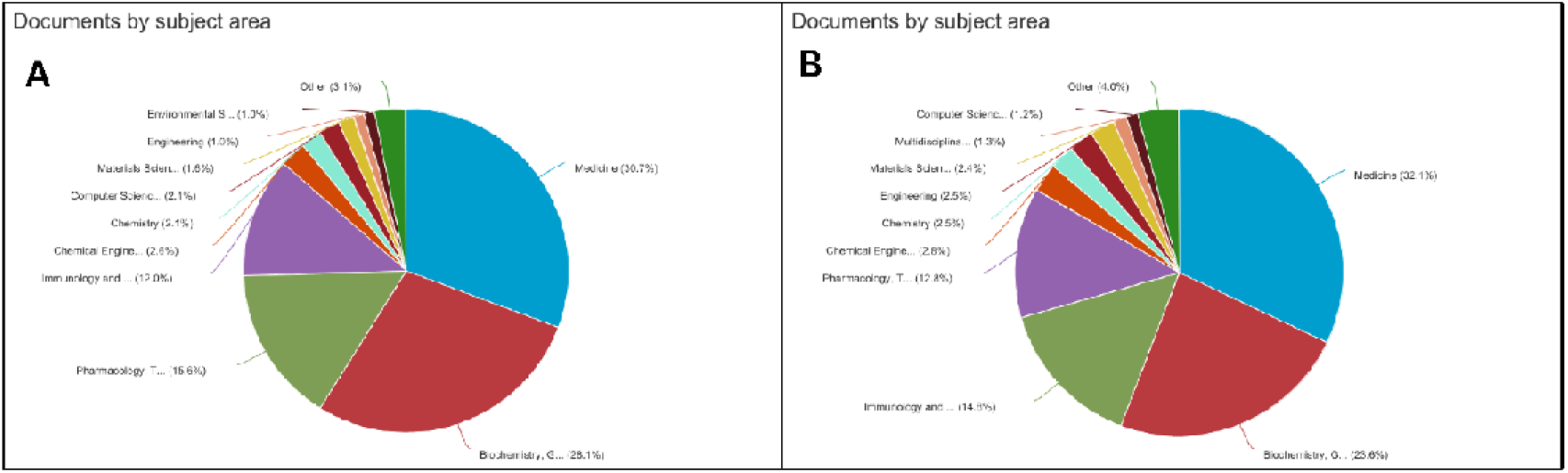
Major subject areas of research A: Circular RNA based cancer vaccines B: RNA based cancer vaccines

The analysis indicates that circRNA-based vaccine research is currently concentrated in core biomedical areas, particularly Medicine and Biochemistry, reflecting its early stage of development. In contrast, RNA-based vaccine research has advanced to include a wider range of interdisciplinary fields, such as Immunology, Microbiology, and materials science, indicative of its more established status.

CircRNA-based vaccine research shows limited engagement in Immunology and interdisciplinary areas like Chemical Engineering and Materials Science. This suggests that as circRNA vaccines advance beyond preclinical stages, there will be a need for increased focus on these areas to address delivery and stability challenges. The current distribution suggests that while circRNA-based vaccines are in an exploratory phase, their growth potential lies in expanding into these additional fields. The relatively low representation in Environmental Sciences and multidisciplinary studies for circRNA vaccines compared to RNA-based vaccines further highlights areas for future expansion. As the field matures, broader interdisciplinary collaborations are expected, similar to those seen in RNA vaccine research.circRNA-based vaccine research is in its nascent stage, with significant potential for growth into interdisciplinary areas. As research progresses, integrating fields like Immunology, Engineering, and Materials Science will be crucial for overcoming development challenges and advancing circRNA vaccines towards clinical application.

### 3.2. In-Depth Focus: Advancing circRNA Vaccines for Cancer

#### 3.2.1. Keywords

The keyword network analysis of circRNA-based vaccines for cancer reveals several key research themes (Figure 7). The central cluster around terms like “circRNA,” “immunotherapy,” and “vaccine” indicates a growing interest in combining circRNAs with cancer immunotherapy. The green cluster emphasizes the use of circRNAs in cancer vaccines and immune stimulation, including their role in “glycolysis” and “chemoresistance.” The red cluster focuses on “circular RNA vaccine,” “mRNA vaccine,” and “cancer therapy,” highlighting a comparative interest in circRNAs and mRNA for vaccine development. The yellow and blue clusters center on “exosome,” “biomarker,” and “small extracellular vesicles,” reflecting interest in circRNAs for vaccine delivery and diagnostics. The orange cluster points to a broader application of circRNA technology in “antiviral therapy.”

**Figure 7.**
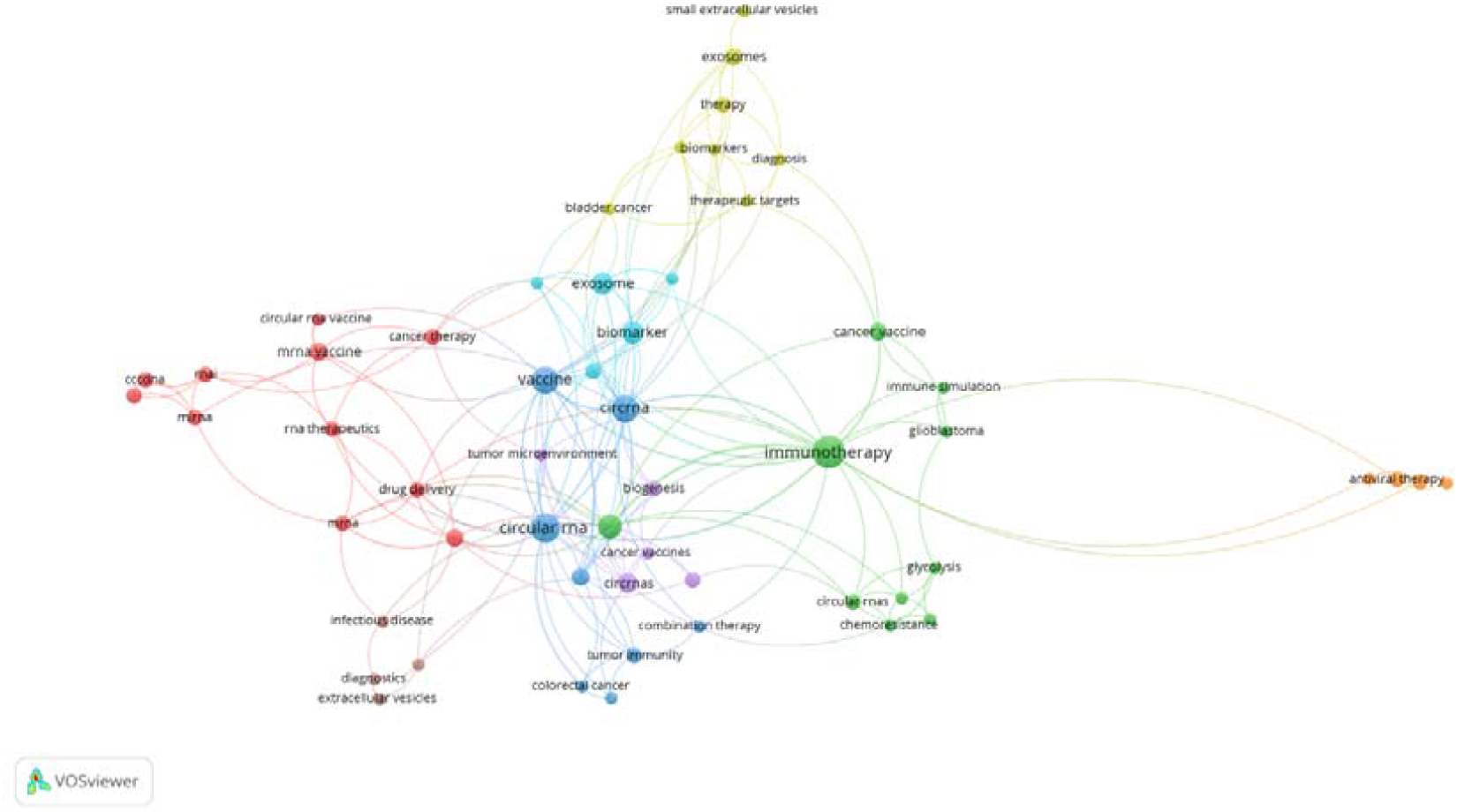
Visualization of keywords network on research of circular RNA vaccines for cancer.

The network analysis illustrates that circRNA-based vaccine research is multifaceted and expanding. The centrality of “circRNA” and “immunotherapy” underscores a significant focus on integrating circRNAs into cancer immunotherapy strategies. The green cluster’s emphasis on immune stimulation and metabolic pathways suggests active exploration of circRNAs in novel therapeutic contexts. The red cluster’s focus on the interplay between circRNAs and mRNA in vaccine development highlights an interdisciplinary approach. The yellow and blue clusters show that circRNAs are being explored for delivery methods and as biomarkers, while the orange cluster indicates potential applications beyond cancer, such as antiviral therapy. These trends reveal a dynamic and evolving research landscape with promising future directions for circRNA-based vaccines in cancer and other medical fields.

#### 3.2.2. Network of Top Cited Authors

The citation network analysis reveals two main clusters of researchers in circRNA-based vaccines for cancer (Figure 8). The first, centered around “Lin, Lin,” includes closely associated authors like “Xu, Yan,” “Sun, Xiaoyu,” and “Chen, Ziqiang,” indicating a dense collaborative network focusing on circRNA mechanisms in cancer vaccines. The second cluster, led by “Amaya, Laura,” is more isolated, suggesting a distinct or emerging line of research. The sparse linkage between clusters indicates potential for future integration and collaboration.

**Figure 8.**
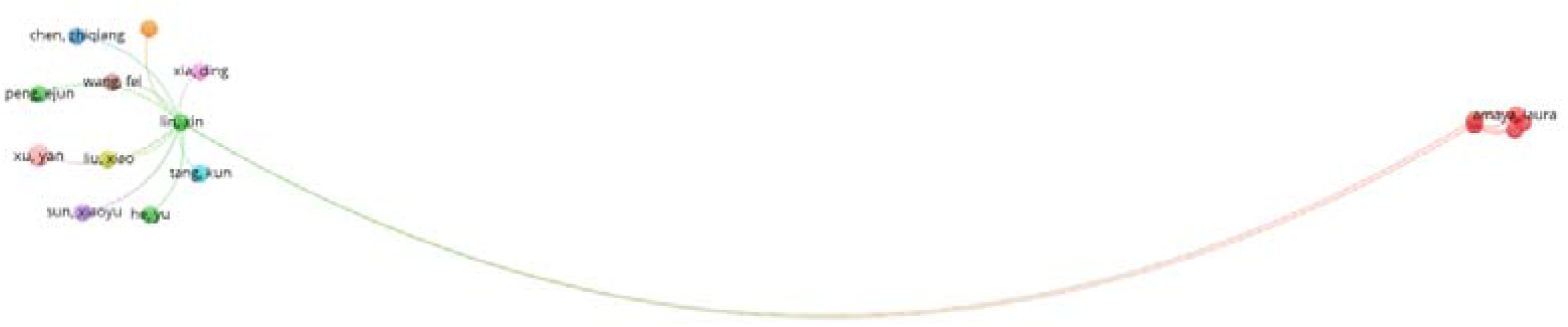
Visualization of top cited authors on research of circular RNA vaccines for cancer.

The analysis shows that while “Lin, Lin” and their network represent a concentrated area of circRNA research, “Amaya, Laura” explores potentially novel aspects of the field. The current separation between clusters reflects diverse research trajectories, but the sparse connections hint at possible future integration. As circRNA research evolves, stronger connections between these clusters may enhance collaboration and unify research efforts, driving significant advancements in cancer vaccine development.

#### 3.2.3. Co-Citation and Bibliometric Coupling Analysis

##### 3.2.1. Authors

As depicted in figure 9, panel A’s co-citation network analysis reveals a dense cluster around “Zhang Y” and “Weissman D,” indicating their central role in circRNA-based vaccine research. Other significant clusters led by “Hansen T.B.” and “Yang L.” show additional influential research groups. Panel B’s bibliometric coupling analysis identifies two distinct clusters: one around “Zhang Y” and “Weissman D,” and another isolated group linked to “Arbuthnot P.” The strong coupling in these clusters suggests cohesive research efforts, while the isolation of “Arbuthnot P.” points to a potentially niche or emerging research direction.

**Figure 9.**
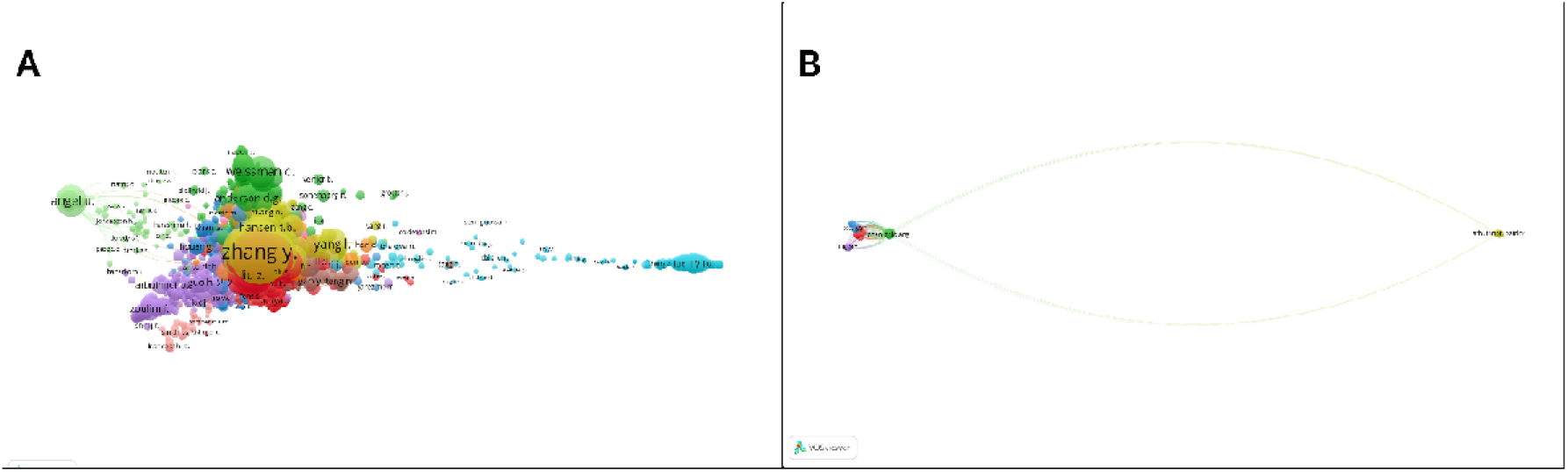
A: Co-citation network analysis and B: Bibliometric coupling analysis of authors doing research on circular RNA cancer vaccines

The co-citation network in Panel A underscores the collaborative nature of circRNA-based vaccine research, with key authors like “Zhang Y” and “Weissman D” being central to foundational work. The bibliometric coupling in Panel B reveals both established research hubs and emerging areas, with “Arbuthnot P.” indicating a new or specialized focus. The integration of these emerging areas with existing research could lead to significant advancements in circRNA-based cancer vaccines, potentially offering novel therapeutic approaches and enhancing patient outcomes.

##### 3.2.3.2. Journal Sources

In the focus of journals in circRNA based cancer vaccine research (Figure 10), panel A’s co-citation network shows “Nature,” “Molecular Cell,” and “Nucleic Acids Research” as central journals, reflecting their key role in circRNA-based vaccine research. Clusters around “Hepatology” and “Oncotarget” indicate cross-disciplinary research linking circRNA with liver diseases and oncology. Panel B’s bibliometric coupling reveals a prominent cluster of journals like “Vaccines,” “Advanced Drug Delivery Reviews,” and “Cancer Letters,” suggesting a strong focus on integrating circRNA into therapeutic applications, particularly in vaccines and drug delivery. Specialized journals such as “Frontiers in Immunology” and “Pharmaceutics” highlight advancements in immunological and formulation aspects.

**Figure 10.**
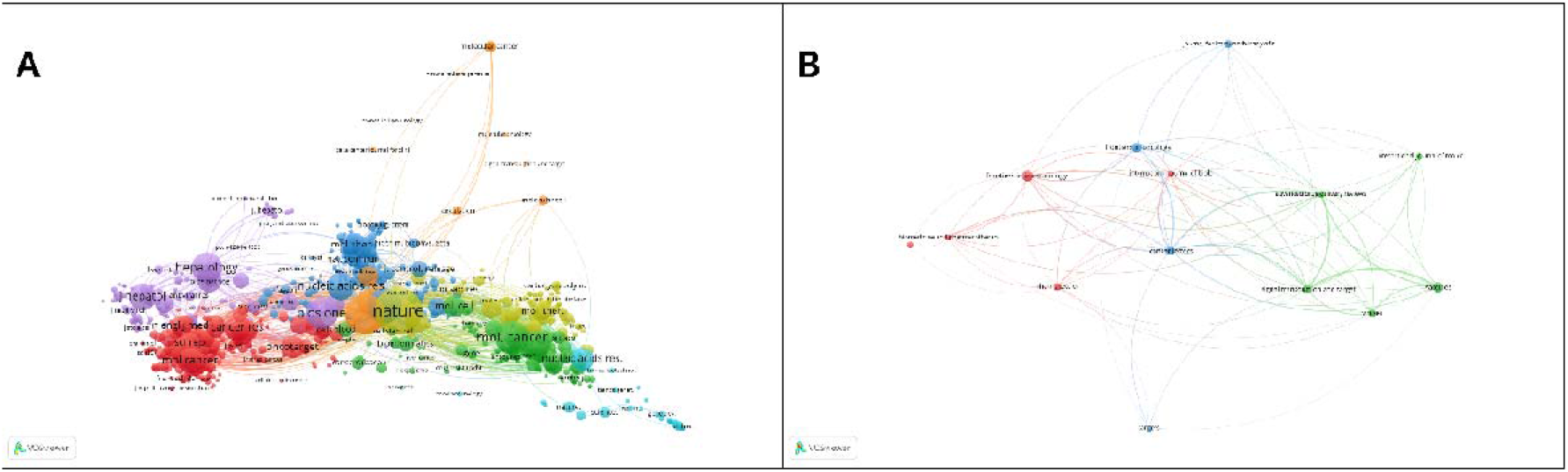
Co-citation network analysis and B: Bibliometric coupling analysis of journals publishing articles on circular RNA cancer vaccines

The co-citation network in Panel A emphasizes the foundational impact of journals like “Nature” and “Molecular Cell” in circRNA-based vaccine research, while also showing significant interdisciplinary connections with journals in liver diseases and oncology. The bibliometric coupling in Panel B underscores the close thematic alignment among journals focused on vaccine development and drug delivery, highlighting the practical integration of circRNA technology. Future growth areas include enhancing circRNA’s role in immunotherapy and optimizing delivery mechanisms, with interdisciplinary journals likely to remain pivotal in advancing these research directions.

##### 3.2.3.3. Organizations

As illustrated in figure 11, panel A’s co-citation analysis reveals a sparse network with institutions like the “Medical Research Center, Sun Y” and the “Department of Urology, Tongji” as key contributors, indicating their significant impact on circRNA-based vaccine research. The “Hunan Provincial Tumor Hospital” also plays a role in niche areas. The limited number of prominent institutions suggests that circRNA research in cancer is still developing. Panel B’s bibliometric coupling shows emerging collaborations among these institutions, such as the “Department of Urology, Tongji” and the “Medical Research Center, Sun Y,” but also indicates a fragmented research landscape with potential for increased integration.

**Figure 11.**
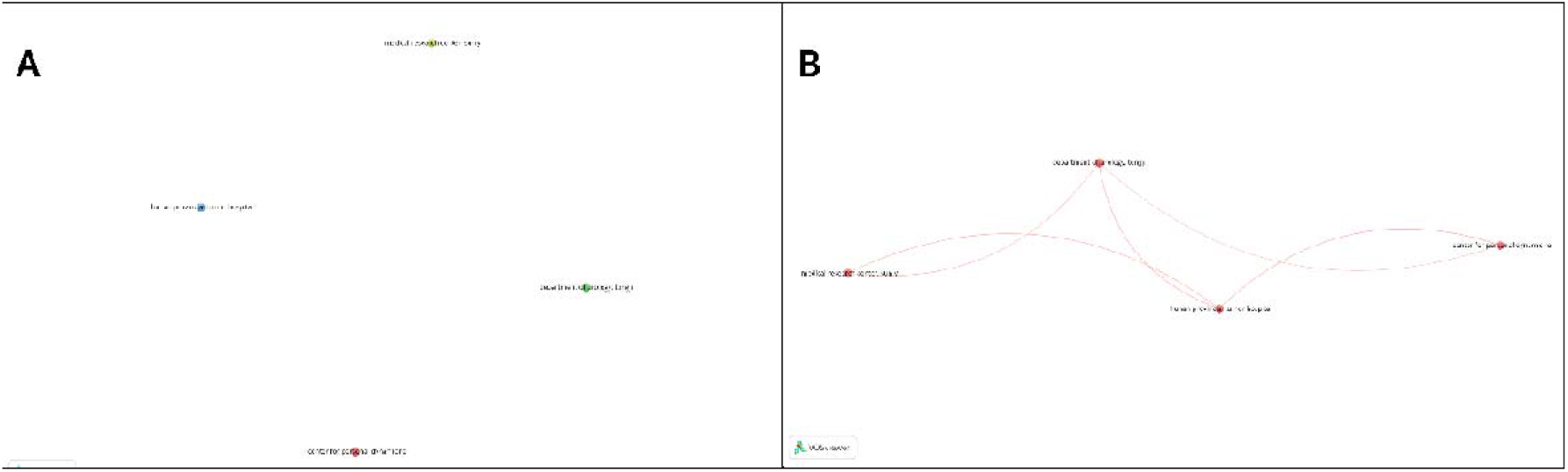
Co-citation network analysis and B: Bibliometric coupling analysis of organizations doing research on circular RNA cancer vaccines

The organizational landscape of circRNA-based vaccine research is nascent, with a few key institutions leading the field. The sparse network in Panel A and the fragmented connections in Panel B highlight both the established contributions of these institutions and the significant opportunities for growth. Enhancing collaboration and encouraging new institutions to join the field could strengthen the research network and accelerate the development of circRNA-based cancer vaccines. As the field evolves, increased interdisciplinary and cross-institutional collaboration is expected to drive further innovation and discoveries.

##### 3.2.3.4. Countries

Shown in figure 12, panel A’s co-citation analysis highlights China and the United States as the leading contributors to circRNA-based vaccine research, with China being the most central and influential. The connections with other countries like South Korea, Germany, and France show China’s prominent role and collaborative efforts. The United States, while impactful, is more isolated in comparison. Panel B’s bibliometric coupling underscores China’s extensive international collaborations and its connections with countries like the United States, Germany, and India. The United States also maintains strong links with Canada, the UK, and Germany. Emerging nations like India, Egypt, and Saudi Arabia are beginning to play more significant roles in the field.

**Figure 12.**
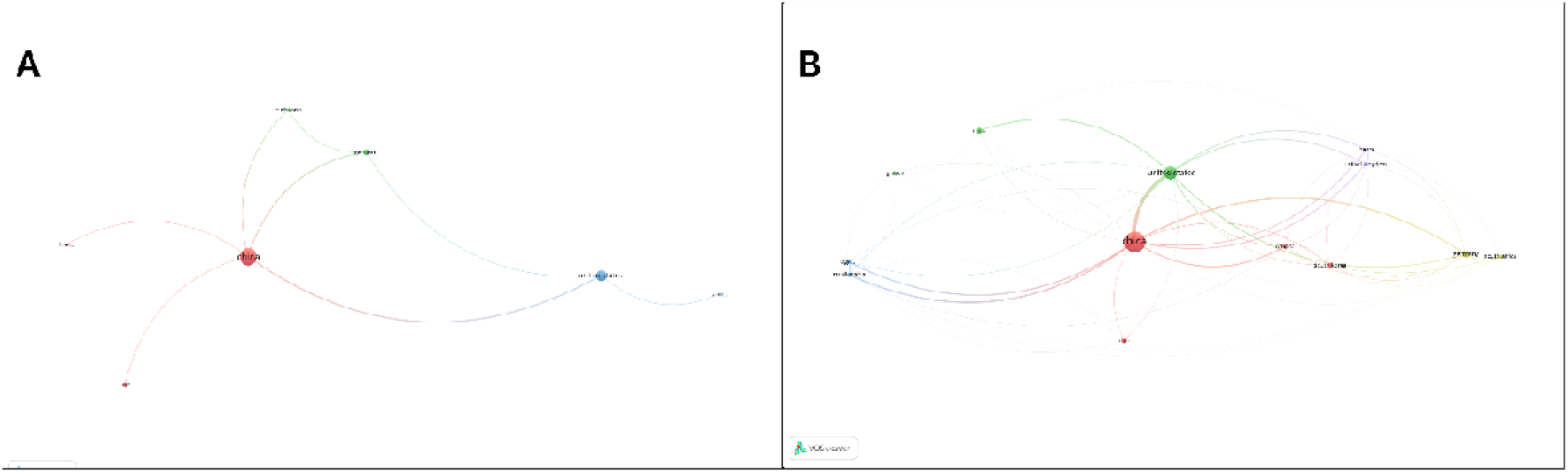
Co-citation network analysis and B: Bibliometric coupling analysis of countries doing research on circular RNA cancer vaccines

The global research landscape for circRNA-based vaccines is currently dominated by China and the United States, with China leading in centrality and collaborative efforts. The United States also plays a key role but is somewhat less collaborative. Emerging countries like Germany, South Korea, and India are increasingly contributing to the field, signaling expanding global engagement. Future advancements are likely to benefit from enhanced international partnerships, integrating new contributors into the research network. Strengthening collaborations between established leaders and emerging nations will be crucial for advancing circRNA-based cancer vaccines and fostering innovative treatments.

## 4.. CONCLUSION

The present study is, to the best of our knowledge, the first to illustrate the current research status and global emerging trends in circular RNA based cancer vaccines research using a bibliometric approach. The global landscape of this rapidly evolving field highlights key trends, with China and the United States emerging as dominant contributors. The study also highlights a shift in research focus from phenotypic studies of cancer towards more mechanistic and therapeutic approaches. Significant research efforts are focused on enhancing circRNA delivery and efficacy, particularly through nanotechnology. Emerging research hotspots, such as the integration of circRNAs with other therapeutic modalities, indicate the potential for multi-modal, personalized cancer therapies. However, the study also identifies the need for more robust clinical trials and expanded international collaborations to fully realize the potential of circRNA-based vaccines in the clinic.

